# Fecal Metabolite Quantitation for Rapid Assessment of the Gut Microbiome

**DOI:** 10.1101/2024.12.13.628394

**Authors:** Michael W. Mullowney, Angelica Moran, Antonio Hernandez, Mary McMillin, Amber Rose, David Moran, Jessica Little, Ann B. Nguyen, Bhakti K. Patel, Christopher J. Lehmann, Matthew A. Odenwald, Eric G. Pamer, Kiang-Teck J. Yeo, Ashley M. Sidebottom

## Abstract

The intestinal microbiome is composed of myriad microbial species that produce metabolites that impact host health. While loss of bacterial species and beneficial metabolites from the fecal microbiome is associated with development of a range of diseases and medical complications, there are currently no diagnostic tests that rapidly identify individuals with microbiome deficiencies. Reduced concentrations of fecal butyrate and deoxycholic acid are associated with adverse clinical outcomes and result from the loss of a subset of health-associated bacterial species. We present a rapid diagnostic test based on 3-nitrophenylhydrazine derivatization and ultrahigh-performance liquid chromatography-mass spectrometry that measures fecal butyrate and deoxycholic acid concentrations as markers of microbiome function. A matrix-matched calibration curve was developed using a simulated fecal mixture to optimize accuracy and facilitate adherence to clinical laboratory regulations. The assay showed an analytical measurement range from 4.3–3030.1 µM (LLOQ = 9.75 µM) for butyrate and from 0.9– 64.9 µM (LLOQ = 0.9 µM) for deoxycholic acid. Precision demonstrated a coefficient of variation <15% at all QC levels tested. The assay can be performed in under an hour from extraction to provision of quantitative results, enabling the rapid identification of patients with defective microbiome function.

## INTRODUCTION

The intestinal microbiome is composed of microbes that provide resistance against infection by enhancing epithelial barrier function and optimizing immune defenses. Commonly prescribed antibiotics can reduce the microbiome’s microbial diversity and low-fiber diets limit microbiome functions, including production of health-promoting metabolites. Loss of microbiome diversity and microbially-produced metabolites has been associated with *Clostridioides difficile* infections^2^, mortality following COVID-19^6^, graft-versus-host disease following hematopoietic stem cell transplantation^1^ and systemic infections in patients with liver disease or following organ transplantation^3,4,5^. Clinical trials investigating the reconstitution of microbiome deficiencies are limited, in part because real-time identification of patients with compositional or functional microbiome deficiencies remains an unmet challenge. Fecal nucleic acid sequencing and metagenomic analyses can determine microbiome compositions and mass spectrometry can quantify fecal metabolites, but these platforms have yet to be developed into clinical tests that rapidly identify patients with deficient microbiome compositions or functions^8^.

While the intestinal microbiome produces thousands of distinct metabolites, short-chain fatty acids (SCFAs) and bile acids are particularly impactful on host physiology and immune defenses. The SCFA butyrate is produced by obligate anaerobic bacterial species in the lower gastrointestinal tract and, in experimental models, impacts immune system development. Primary bile acids are produced and conjugated to amino acids in the liver and, upon delivery to the gut, facilitate nutrient absorption. Intestinal microbes, however, modify bile acids by deconjugating amino acids from the bile acid core and convert primary to secondary bile acids by modifying bile acid hydroxylation states. Secondary bile acids, such as deoxycholic acid (DCA) and its microbially-generated derivatives, impact host physiology and immune cell differentiation. Butyrate and DCA concentrations are commonly reduced in hospitalized patients, generally a result of antibiotic-mediated loss of microbiome microbial diversity and are associated with a range of adverse medical outcomes.

Given the dynamic nature of the intestinal microbiome compositions, real time insight into the risk of adverse health outcomes will require rapid quantitation of fecal metabolite concentrations. Clinical certification by CLIA and other regulatory authorities requires a matrix-matched calibration curve for more accurate measurement of metabolites.^13^ While simulated fecal matrices are being developed for metagenomic sequencing, matrices specific for targeted metabolomics are currently unavailable. Herein, we developed a simulated fecal matrix to measure the microbiota-derived metabolites. With the goal of same day identification of patients with reduced intestinal microbiome function, we have optimized the method to measure fecal butyrate and deoxycholic acid (DCA) on the same LC-MS platform. The method has been developed, optimized, and validated as a potential platform to screen candidates for clinical trials of microbiome reconstitution.

## MATERIALS AND METHODS

### Materials

A solution of 80% LC-MS grade methanol (Fisher Scientific; Optima A456) with D_3_-acetate (Cambridge Isotope Laboratories; DLM-3126), D_5_-propionate (Cambridge Isotope Laboratories; DLM-1601), 500 µM sodium D_7_-butyrate (Cambridge Isotope Laboratories; DLM-7616), D_9_-valerate (Cambridge Isotope Laboratories; DLM-572), D_4_-cholate (Cambridge Isotope Laboratories; DLM-2611), 2.55 µM D_4_-deoxycholic acid (Cambridge Isotope Laboratories; DLM-2824), D_4_-glycocholate (Cambridge Isotope Laboratories; DLM-2742), D_4_-lithocholate (Cambridge Isotope Laboratories; DLM-9560), and D_4_-taurocholate (Cambridge Isotope Laboratories; DLM-9572) as internal standards (ISTDs) was prepared for extraction of all samples.

LC-MS/MS methods were optimized and calibration curves were built using authenticated standards of acetate (Sigma Aldrich; S2889), propionate (Sigma Aldrich; P1880), butyrate (Sigma Aldrich; B5887), isobutyrate (Sigma Aldrich; I1754), valerate (Sigma Aldrich; 240370), desaminotyrosine (Sigma Aldrich; H52406), indole-3-carboxaldehyde (Sigma Aldrich; 129445), cholate (Sigma Aldrich; C1129), glycocholate (Sigma Aldrich; G2878), deoxycholic acid (Sigma Aldrich; D2510), ursodeoxycholic acid (UDCA; Sigma Aldrich; U5127), hyodeoxycholic acid (HDCA; Sigma Aldrich; H3878), isodeoxycholic acid (iso-DCA; 3-epideoxycholic acid; Steraloids, Inc.; C1165-000), chenodeoxycholic acid (CDCA; Sigma Aldrich; C9377), 3-deoxycholic acid (3-DCA; Steraloids, Inc.; C1170-000), taurocholate (Sigma Aldrich; T4009), tauro-alpha-muricholate (Avanti Polar Lipids; 700243P), tauro-beta-muricholate (Steraloid, Inc.; C1899-000), isoallolithocholate (Avanti Polar Lipids; 700196), isolithocholate (Avanti Polar Lipids; 700195P), and lithocholate (Sigma Aldrich; L6250).

Derivatization reagents were *N*-(3-dimethylaminopropyl)-*N*′-ethylcarbodiimide hydrochloride (EDC; Sigma-Aldrich; E7750) and 3-nitrophenylhydrazine hydrochloride (3-NPH; Sigma-Aldrich; N21804). Solvents used were LC-MS grade water (Fisher Scientific; Optima W7), LC-MS grade acetonitrile (Fisher Scientific; Optima A955), LC-MS grade 2-propanol (Fisher Scientific; Optima A461), and 99% Ultra-Pure LC-MS Grade formic acid (CovaChem; 11202) as a mobile phase buffer.

The simulated fecal matrix was comprised of non-organic avocado (Trader Joe’s), Gluten-Free All Purpose Flour (comprised of millet flour, potato starch, tapioca starch, rice flour, sorghum flour, and xanthan gum; Trader Joe’s), canned 365 Whole Foods Market organic black beans (Whole Foods Market), Clabber Girl Cornstarch (Jewel-Osco Grocery), and Xclusiv Organics cosmetic grade snail mucilage extract powder (Walmart.com).

### Simulated fecal matrix

Two microbiota-derived metabolites, butyrate and deoxycholic acid, were quantified in food products and snail mucin using LC-MS/MS. Food items with endogenous concentrations of these metabolites below the limit of quantitation were selected to create a simulated fecal matrix with a consistency similar to human stool. The simulated fecal mixture consisted of avocado, gluten-free all-purpose flour, black beans, corn starch, and snail mucilage. These solid components were blended until smooth in a conventional household blender and diluted with 0.7 mL deionized Milli-Q water per gram of solid mixture.

### Clinical sample collection

Approval by the University of Chicago Institutional Review Board was granted for the clinical studies that were the source of the samples analyzed herein (liver disease, IRB21-0327; liver transplant, IRB 20-0163; heart transplant, IRB20-0333; medical intensive care unit, IRB20-1102).

### Sample extraction and preparation

Extraction solvent (80% methanol spiked with internal standards purchased from Cambridge Isotope Laboratories) was added to all pre-weighed samples to 100 mg/mL in beadruptor tubes (Fisherbrand; 15-340-154). Samples were homogenized at 4 ^°^C on a Bead Mill 24 Homogenizer (Fisher; 15-340-163), set at 1.6 m/s with 6 thirty-second cycles, 5 seconds off per cycle. Samples were then centrifuged at −10 ^°^C, 20,000 x *g* for 15 min and the supernatant was used for subsequent metabolomic analysis.

### Conventional Metabolite Analysis using GC-MS with PFB-Br Derivatization

Short-chain fatty acids were derivatized as described by Haak, *et al*. with the following modifications.^14^ Metabolite extract (100 µL) was added to borate buffer (100 µL) (pH 10) (Thermo Fisher, 28341), 400 µL of 100 mM pentafluorobenzyl bromide (Millipore Sigma; 90257) in acetonitrile, and 400 µL of *n*-hexane (Acros Organics; 160780010) in a capped MS autosampler vial (Microliter; 09-1200). Samples were heated in a thermomixer C (Eppendorf) to 65 ^°^C for 1 hour while shaking at 1300 rpm. After cooling to room temperature, samples were centrifuged at 4 ^°^C, 2000 x *g* for 5 min, allowing phase separation. From the hexanes phase (top layer), 100 µL was transferred to an autosampler vial containing a glass insert and the vial was sealed. Another 100 µL of the hexanes phase was diluted with 900 µL of *n*-hexane in an autosampler vial. Concentrated and dilute samples were analyzed using a GC-MS (Agilent 7890A GC system, Agilent 5975C MS detector) operating in negative chemical ionization mode, using a HP-5MSUI column (30 m x 0.25 mm, 0.25 µm; Agilent Technologies 19091S-433UI), methane as the reagent gas (99.999% pure) and 1 µL split injection (1:10 split ratio). Oven ramp parameters: 1 min hold at 60 ^°^C, 25 ^°^C per min up to 300 ^°^C with a 2.5 min hold at 300 ^°^C. Inlet temperature was 280 ^°^C and transfer line was 310 ^°^C. A 10-point calibration curve was prepared with acetate (100 mM), propionate (25 mM), butyrate (12.5 mM), and succinate (50 mM), with nine subsequent 2x serial dilutions. Data analysis was performed using MassHunter Quantitative Analysis software (version B.10, Agilent Technologies) and confirmed by comparison to authentic standards. Normalized peak areas were calculated by dividing raw peak areas of targeted analytes by averaged raw peak areas of internal standards.

### Conventional Bile Acid Analysis using QToF LC-MS

Bile acids were analyzed using LC-MS. A 75 µL aliquot of metabolite extract was added to prelabeled MS autosampler vials (Microliter; 09-1200) and dried down completely under a nitrogen stream at 30 L/min (top) 1 L/min (bottom) at 30 ^°^C (Biotage SPE Dry 96 Dual; 3579M). Samples were resuspended in 750 µL of 50:50 water:methanol. Vials were added to a thermomixer C (Eppendorf) to resuspend analytes at 4 ^°^C, 1000 rpm for 15 min with an infinite hold at 4 ^°^C. Samples were transferred to prelabeled microcentrifuge tubes and centrifuged at 4 ^°^C, 20,000 x *g* for 15 min to remove insoluble debris. A 700 µL portion of supernatant was transferred to a fresh, prelabeled MS vial. Samples were analyzed on an Agilent 1290 infinity II liquid chromatography system coupled to an Agilent 6546 quadrupole time-of-flight (QToF) mass spectrometer operating in negative mode, equipped with an Agilent Jet Stream Electrospray Ionization source.

A sample volume of 5 µL was injected onto an Xbridge© BEH C18 Column (3.5 µm, 2.1 × 100 mm; Waters Corporation, 186003022) fitted with an XBridge© BEH C18 guard (Waters Corporation, 186007766) at 45 ^°^C. Mobile phase A was water with 0.1% formic acid and mobile phase B was acetone with 0.1% formic acid. Gradient elution started with 28% B with a flow rate of 0.4 mL/min for 1 min and linearly increased to 33% B over 5 min, then linearly increased to 65% B over 14 min. Then the flow rate was increased to 0.6 mL/min and B was increased to 98% over 0.5 min. These conditions were held constant for 3.5 min. Re-equilibration was completed at a flow rate of 0.4 mL/min of 28% B for 3 min. The electrospray ionization conditions were set with the capillary voltage at 3.5 kV, nozzle voltage at 2 kV, and detection window set to 100-1700 *m/z* with continuous infusion of a reference mass (Agilent ESI TOF Biopolymer Analysis Reference Mix) for mass calibration.

A ten-point calibration curve was prepared with 125 µg/mL each in water of cholic acid, taurocholic acid, glycocholic acid, deoxycholic acid, 3-deoxycholic acid, alloisolithocholic acid, and lithocholic acid with nine subsequent 3x serial dilutions. Data analysis was performed using MassHunter Profinder Analysis software (version B.10, Agilent Technologies) and confirmed by comparison with authentic standards. Normalized peak areas were calculated by dividing raw peak areas of targeted analytes by averaged raw peak areas of internal standards.

### Rapid metabolomic screen using LC-MS

Based on methods from Liao, *et al.^12^*, 25 µL of samples were transferred to a mass spectrometry vial with 12.5 µL 3-nitrophenylhydrazine (3-NPH) and 12.5 µL N-(3-dimethylaminopropyl)-N′-ethylcarbodiimide hydrochloride (EDC). Samples were incubated at 40 ^°^C for 30 minutes and then chilled at −80 ^°^C for 2 minutes to complete the derivatization process (**Fig 1a**). For optimal separation of isomers, 2 µL of sample was injected and separated on a CORTECS T3 column (Waters, 120Å, 1.6 µm, 2.1 × 100 mm) fitted with a CORTECS T3 VanGuard Pre-column (Waters, 120Å, 1.6 µm, 2.1 mm X 5 mm) at 45^°^C with a flow rate of 0.550 mL/min and run-time of 10 min using a Sciex Exion LC AD system (**Fig 1b**). Mobile phase A was water with 0.1% formic acid and mobile phase B was isopropanol: acetonitrile (3:1) with 0.1% formic acid. Separation began with a 1 min hold at 15% B which was followed by 15–48% B from 1–4 min, 48% B for 2 min, 48–100% B from 6–8 min, 100% B from 8–9 min, 100–15% B from 9–9.1 min, and 15% B from 9.1–10 min. Analytes were detected by negative ionization using a Sciex QTRAP 6500 mass spectrometer with an electrospray ionization source. The method parameters are as follows, declustering potential: −60, entrance potential: −10, collision cell exit potential: −13, curtain gas: 30, collision gas: 12, ion spray voltage: −4500, temperature: 325, ion source gas 1: 40, and ion source gas 2: 40.

**Fig 1.**
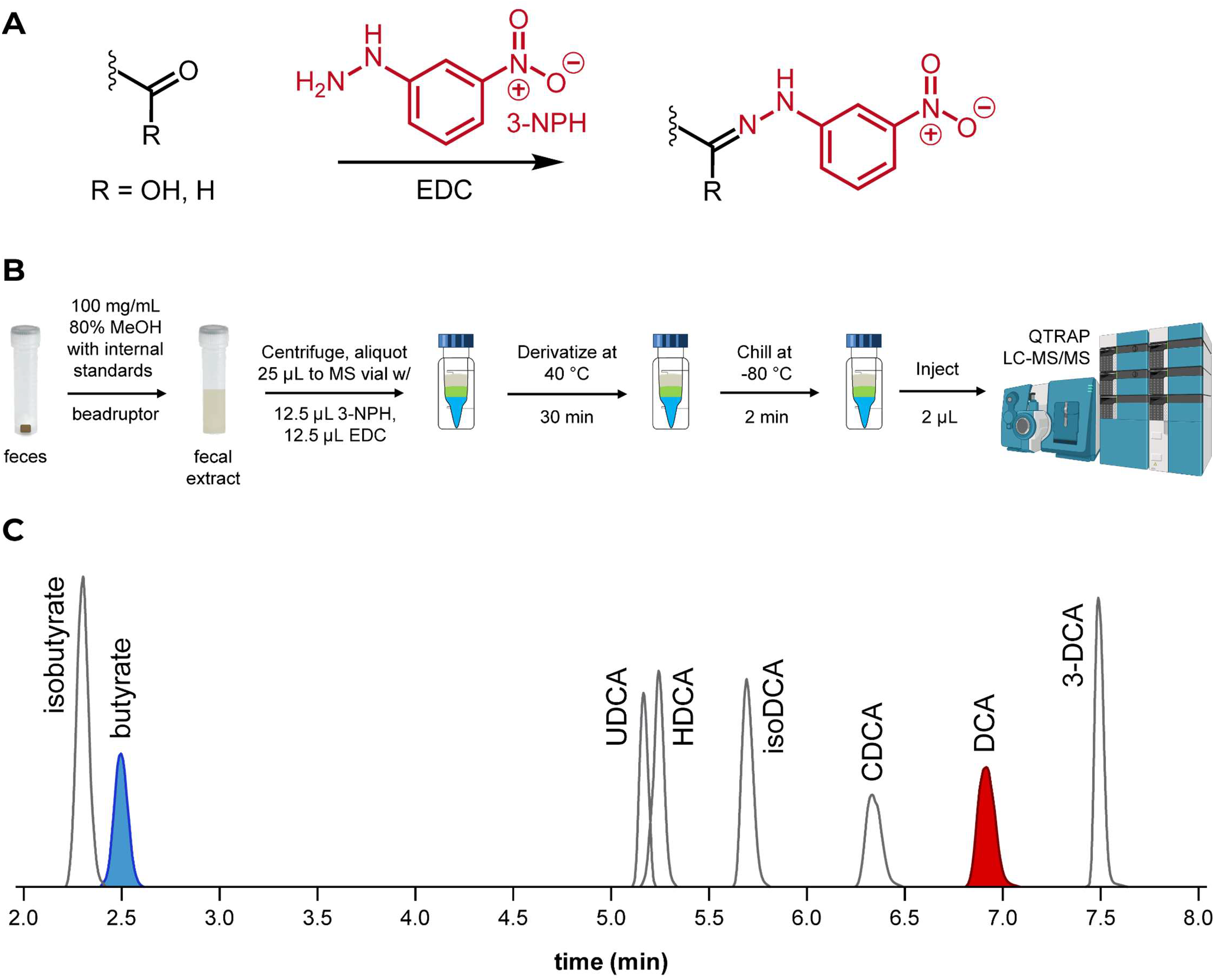
(**A**) Reaction mechanism of carboxylic acid and aldehyde derivatization with 3-nitrophenylhydrazine (3-NPH) catalyzed by *N*-(3-(dimethylamino)propyl)-*N*′-ethylcarbodiimide (EDC). (**B**) Five-step procedure to derivatize and prepare patient fecal samples for LC-MS/MS analysis. (**C**) A 10-minute chromatographic method enables the detection and resolution of butyrate and deoxycholic acid from their isomers in the same sample injection.

Data analysis was performed using MultiQuant™ software (version 3.0, Sciex). The precursor, quantifier product, and qualifier ions and retention times (± 0.2 min) for 3-NPH derivatized metabolites of interest were: 3-NPH-butyrate (precursor: *m/z* 222.09, quantifier: *m/z* 137.1, qualifier: *m/z* 152.00, 2.40 min) and 3-NPH-deoxycholic acid (precursor: *m/z* 526.33, quantifier: *m/z* 152.00, qualifier: *m/z* 137.1, 6.81 min). Derivatized biologically relevant isomers were measured based on detection of the same ions with unique retention times when compared to butyrate (isomer: isobutyrate) and deoxycholic acid (isomers: ursodeoxycholic acid, hyodeoxycholic acid, 3-deoxycholic acid, chenodeoxycholic acid, and isodeoxycholic acid; **Fig 1c**). Acquisition parameters can be found in **Table S2**. For quantification, peak areas of 3-NPH derivatized metabolites in each sample were normalized to the peak areas of their respective internal standards (D_7_-3-NPH-butyrate for butyrate, D_4_-3-NPH-deoxycholic acid for deoxycholic acid).

### Method Validation of Rapid Metabolomic Screen

#### Preparation of Calibrators and Quality Control (QC) Materials

Calibrators and QCs were created by spiking butyrate and deoxycholic acid into either 50% methanol (for conventional metabolite analyses) or simulated fecal matrix (rapid metabolomic screen validation). A set of seven calibrators were prepared for butyrate (50–4000 µM) and deoxycholic acid (1–80 µM). QCs were independently prepared to five concentration levels of butyrate (125, 375, 750, 1250, and 2500 µM) and deoxycholic acid (2.5, 7.5, 15, 25, 50 µM).

#### Analytical Measurement Range (AMR) and the Lower Limit of Quantification (LLOQ) Studies

The AMR was determined by creating a set of simulated fecal matrix samples with butyrate and deoxycholic acid at concentrations ranging from 5–3250 µM and 0.1–65 µM, respectively. The measured concentrations were compared to the expected concentrations to assess linearity. The LLOQ was determined by running a set of 5 low concentration simulated samples over 3 days (8 replicates/day) and defined as concentration at which the % CV across replicates is <20 %.

#### Precision and Recovery Studies

Intra-day and inter-day precision was assessed using QC materials with known amounts of butyrate and deoxycholic acid in the simulated fecal matrix. QCs were measured 10 times in a single run to determine intra-day precision. Inter-day reproducibility was determined by testing QCs in duplicate daily for a total of 14 replicates. CV of <15 % at each level was considered acceptable. Accuracy was assessed by calculating the recovery of each analyte relative to the expected concentrations.

#### Method Comparison

Clinical samples were tested using conventional metabolite analysis by PFB-Br derivatization with GC-MS, conventional bile acid analysis by LC-MS, and the rapid metabolomic screen. Results from samples within the analytical measurement range of all methods were analyzed using linear regression and Bland–Altman bias plots to determine the agreement between methods.

## RESULTS

Short-chain fatty acids are commonly measured by gas chromatography-mass spectrometry (GC-MS) following derivatization with pentafluorobenzyl bromide (PFB-Br) or trimethylsilyl (TMS)-donating reagents such as *N,O*-bis(trimethylsilyl)trifluoroacetamide (BSTFA; **Fig 2a**).^10^ Bile acids, on the other hand, are commonly measured on liquid chromatography-MS (LC-MS) systems using sample preparation that may include steps to optimize signal such as dry down and resuspension, but without derivatization (**Fig 2b**). LC-MS-based platforms allow sample preparation and analysis in significantly less time when compared to GC-MS methods, but SCFAs suffer from poor recovery, stability, and ionization in conventional LC-MS methods.^11^ Conversely, larger metabolites like bile acids lack the volatility for efficient and sensitive GC-based analysis. Among the requirements for a method encompassing both classes of metabolites were the applicability to one LC-MS-based system with one injection, a single calibration curve, a single set of matched internal standards, and quality control samples that accounted for all chemical species. The chemical characteristic between the SCFAs and bile acids is the carboxylic acid functional group. With this in mind, we identified a shorter method involving derivatization of the carboxylic acid moiety by 3-nitrophenylhydrazine (3-NPH) catalyzed by *N*-(3-dimethylaminopropyl)-*N*′-ethylcarbodiimide hydrochloride (EDC),^12^ enabling measurement of both SCFAs and bile acids on the same LC-MS/MS platform (**Fig 2c, d**).

**Fig 2.**
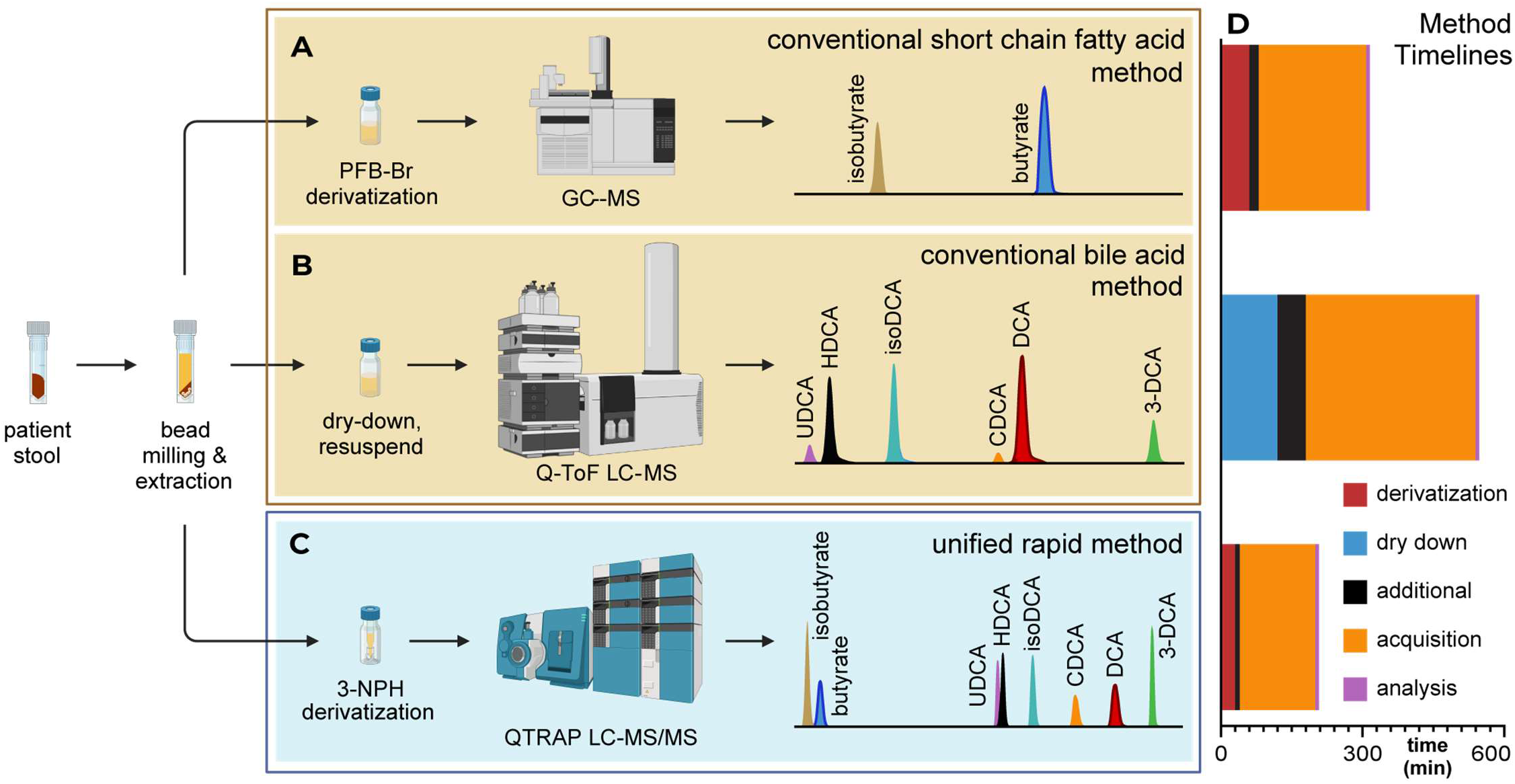
Conventional targeted metabolomics methods require multiple mass spectrometry platforms to measure metabolites with dissimilar chemical characteristics. (**A**) Short-chain fatty acids (SCFAs) such as butyrate typically require analysis by GCMS, often with derivatization, while (**B**) bile acids are routinely analyzed using LC-MS technology. (**C**) The rapid mass spectrometry method presented here unifies the measurement of SCFAs and bile acids while resolving multiple isomers on a single system that minimizes sample preparation and data acquisition time. (**D**) Sample preparation, data acquisition, and analysis time to acquire quantitative butyrate and deoxycholic acid concentration data for one biological sample using conventional metabolomics methods and the rapid metabolomics screen (See also **Table S1**).

Chromatogram overlays from the second lowest calibrator (50 µM) in the matched matrix for butyrate quantifier and qualifier transitions with the D_7_-butyrate internal standard (500 µM) quantifier transition are shown in **Fig 3a** and **b**. Butyrate concentrations in experimental samples were determined using the butyrate quantifier transition of mass-to-charge ratio (*m/z*) 222.1 to 137.1 normalized to the D_7_-butyrate internal standard quantifier mass transition of *m/z* 229.1 to 137.1. Chromatogram overlays from the lowest calibrator (1 µM) in matched matrix for both deoxycholic acid quantifier and qualifier transitions with the D_4_-deoxycholic acid internal standard (2.5 µM) quantifier transition are shown in **Fig 3c** and **d**. Deoxycholic acid concentrations in experimental samples were determined using the deoxycholic acid quantifier transition of *m/z* 526.3 to 152.0 normalized to the D_4_-deoxycholic acid internal standard quantifier mass transition of *m/z* 530.3 to 137.1.

**Fig 3.**
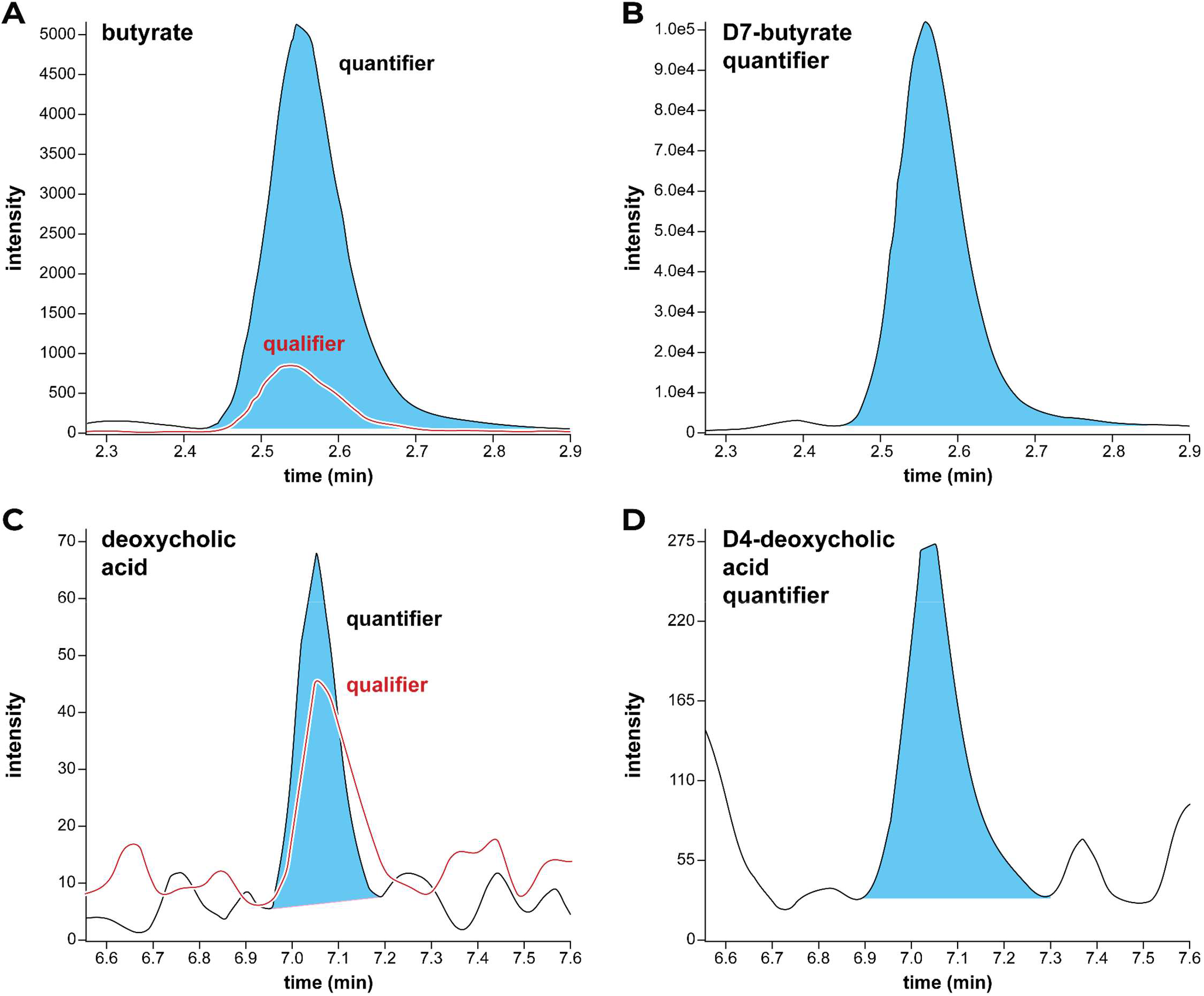
Multiple reaction monitoring chromatogram overlays from the lowest rapid metabolomic screen calibrator for butyrate (**A**; quantifier, *m/z* 222.1–137.1; qualifier, *m/z* 222.1–152.0) with D_7_-butyrate internal standard quantifier transition (**B**; *m/z* 229.1–137.1) and deoxycholic acid (**C**; quantifier, *m/z* 526.3–152.0; qualifier, *m/z* 526.3–137.1) with D_4_-deoxycholic acid internal standard quantifier transition (**D**; *m/z* 530.3–137.1).

### AMR and LLOQ

The rapid metabolomic screen demonstrates an AMR from a butyrate concentration of 4.3–3030.1 µM with an r^2^ value of 0.9991 and from a deoxycholic acid concentration of 0.9–64.9 µM with an r^2^ value of 0.9980 (**Fig 4a, b**). Using a % CV cutoff of 20 %, the lower limit of quantitation (LLOQ) was determined to be 9.75 µM for butyrate and 0.9 µM for deoxycholic acid (**Fig 4c, d**).

**Fig 4.**
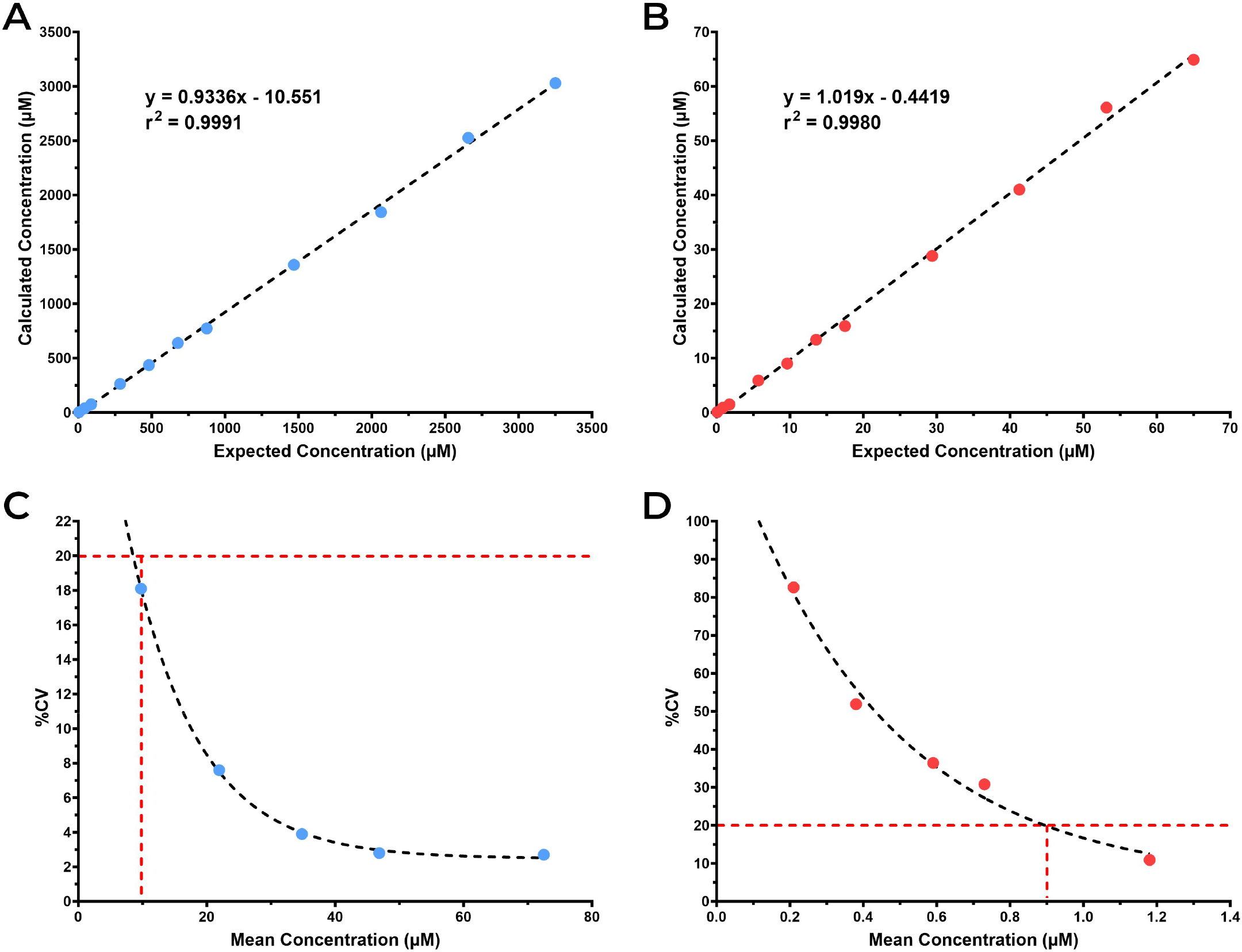
The analytical measurement ranges (AMR) and linearity of butyrate (**A**) and DCA (**B**) are shown across eleven concentration points for the rapid metabolomic screen. The limits of quantitation (LoQ) for butyrate (**C**) and deoxycholic acid (**D**) are defined by the concentration at which the percent coefficient of variation (% CV) reaches 20 %.

### Precision and Accuracy

The reproducibility of this assay is excellent, with an inter-day precision of <15 % CV at all five levels tested across the analytical measurement range (**Table 1)**. Recovery ranged from 98.2–109% for butyrate and 92.4– 109.7 % for deoxycholic acid, demonstrating accuracy of analyte measurement relative to expected concentrations using simulated fecal samples. Additional performance measures of the rapid metabolomic screen can be found in **Table S3**.

**Table 1.**
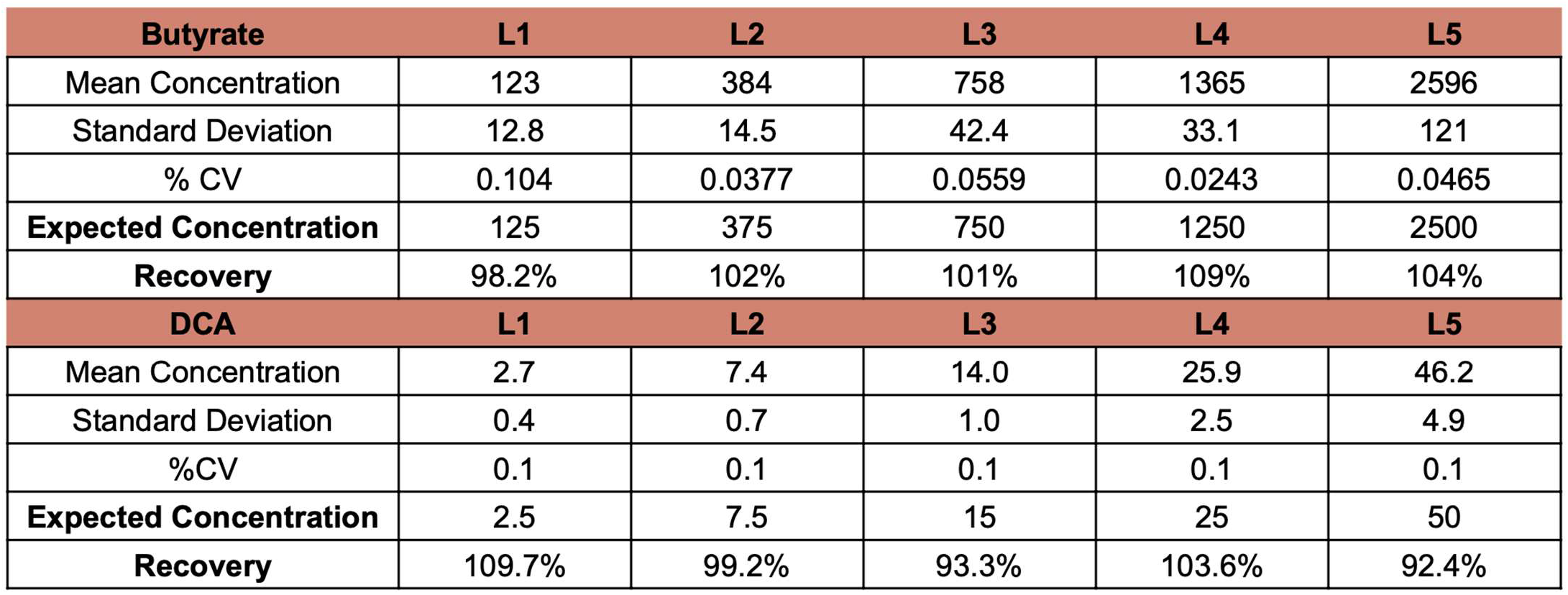
Inter-day precision and accuracy for five concentration levels (L1–L5) spanning the analytical measurement range of the rapid metabolomic screen. Concentrations are shown in μM; % CV = percent coefficient of variation.

### Method Comparison

A total 78 patient fecal samples from across MICU, heart transplant, liver disease, and liver transplant cohorts were used for method comparison. Fecal samples were tested using conventional metabolite analysis by GC-MS with PFB-Br derivatization, conventional bile acid analysis by LC-MS, and the rapid metabolomic screen. Within this set, there were 33 patients with butyrate concentrations within the AMR of both the rapid metabolomic screen and our conventional GC-MS with PFB-Br method, while there were 30 patients with deoxycholic acid concentrations within the AMR of both the rapid metabolomic screen and the conventional QToF LC-MS-based method. For each patient, the butyrate concentration measured by the rapid metabolomic screen and conventional GC-MS analysis were compared for linearity and bias (**Figs 5a, b**) and showed an overall bias of 313 µM (Bland-Altman analysis). A comparison of the DCA concentrations measured using the rapid metabolomic screen and the conventional bile acid LC-MS analysis to assess linearity and bias showed a bias of −3.3 µM (**Figs 5c, d**).

**Fig 5.**
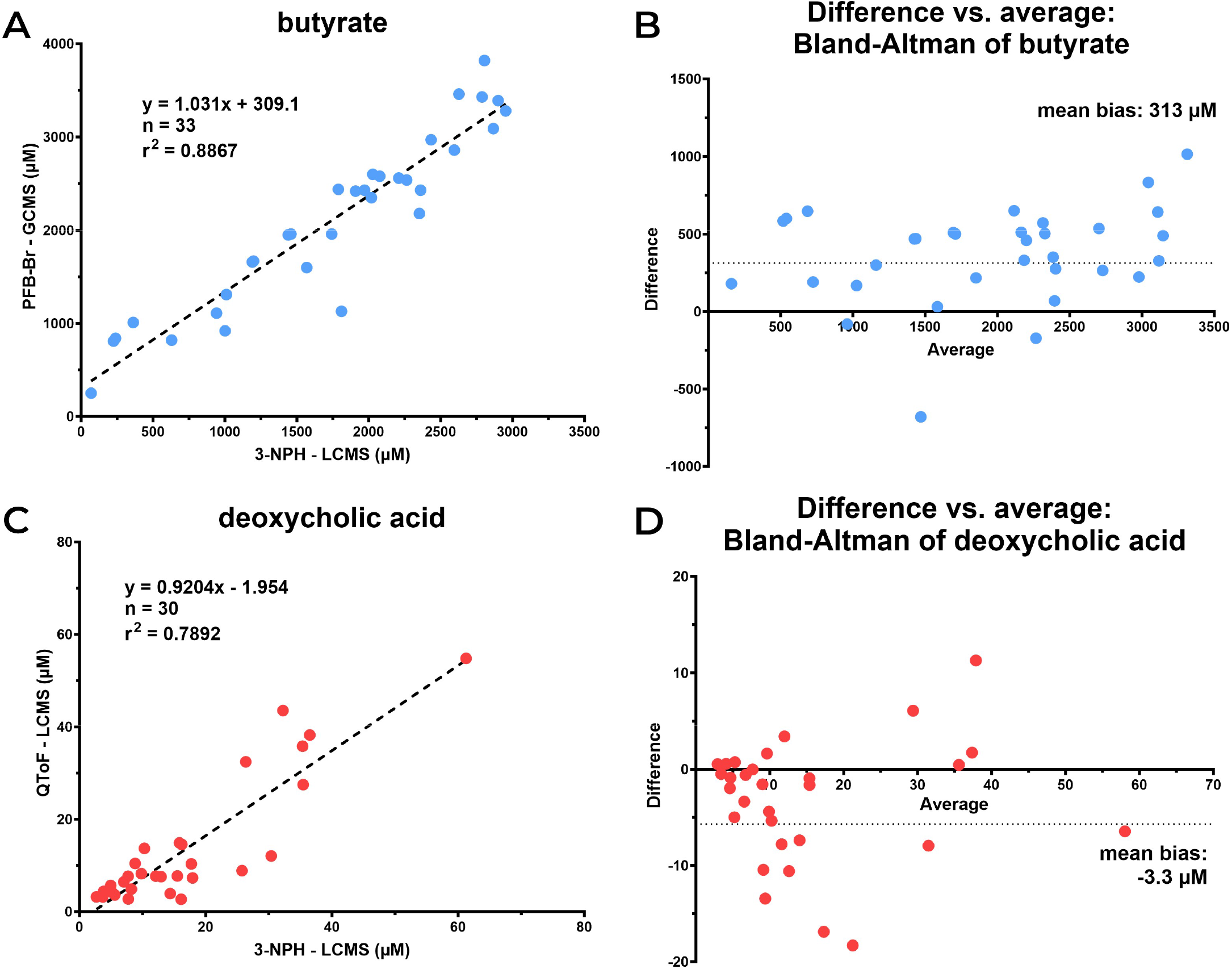
Comparison of the rapid metabolomic screen versus measurement by conventional mass spectrometry methods in patient fecal samples. Linear regression (**A**) and Bland-Altman (**B**) comparisons of butyrate concentrations as measured by the rapid metabolomic screen versus measurement by conventional GC-MS metabolite analysis with PFB-Br derivatization (GCMS); Linear regression (**C**) and Bland-Altman (**D**) comparisons of deoxycholic acid concentrations as measured by the rapid metabolomic screen versus measurement by conventional bile acid analysis using QToF LC-MS. See **Table S4** for patient butyrate and deoxycholic acid concentrations with clinical metadata.

## DISCUSSION

We have developed a rapid metabolomic screen to identify patients with microbiome insufficiency. Fecal microbiome metabolomics has promise in evaluating patient health, but current methods are time-intensive and not widely implemented. The chemical characteristics of different target metabolites often require specific and differing sample preparation and instrumentation platforms for optimal, efficient, and reproducible quantitation. Thus, accurately and expeditiously measuring multiple metabolites in a single screen is problematic because the optimal method for one metabolite is often incompatible with another. The method we developed herein accelerates the quantitation of two key fecal metabolites, butyrate and DCA, by using 3-NPH derivatization and QTRAP quantitation and establishes an LC-MS/MS platform that provides same-day insights into the fecal microbiome’s function.

Until recently, identifying patients with dysfunctional microbiomes was principally of academic interest since the ability to improve microbiome compositions and functions was limited. As the roles of specific microbes and microbiome-derived metabolites in disease resistance have been discovered, however, there is increasing interest in the design, development and conduct of clinical trials to modify, diversify or optimize microbiome compositions and functions. Specifying microbiome deficiencies will be critical for the conduct and success of clinical trials since there are many ways that the complex microbial populations constituting the intestinal microbiome can become insufficient. While the metabolomic screen we describe will rapidly identify patients with defective microbiome function who might be enrolled in clinical trials of microbiome reconstitution, it also represents a step in the direction of personalizing microbiome-targeting interventions by targeting specific defects. We demonstrate that additional microbiome-relevant metabolites can be quantified by the rapid screen, enabling assessment of more complex patterns of microbiome health and facilitating a more personalized approach. In addition to expanding the set of metabolites measured in fecal samples, the test can be adapted for other biological sample types, such as plasma or urine, which are more readily acquired from patients.

A limitation of the method is in its inability to distinguish between patients with low concentrations of butyrate attributable to depleted microbiome diversity versus low prebiotic fiber intake or other causes unrelated to microbiome composition. The addition of a quantitative PCR method targeted for specific organisms may be a useful adjunct to rapidly assess for compositional disturbances.

We observed differences in the quantitation of butyrate and DCA between the rapid screen and the conventional platform. There is a positive bias when comparing 3-NPH derivatization and QTRAP measurement of butyrate in the rapid metabolomic screen to our conventional measurements using PFB-Br derivatization and GC-MS. While differing reaction efficiencies between 3-NPH and PFB-Br may contribute to these variations, bias could be impacted due to sensitivity differences between the instrument mass analyzers. The QTRAP in the rapid metabolomic screen allows for much higher sensitivity in the detection of butyrate, with an LLOQ at 9.75 µM, compared to an LLOQ of 750 µM in the conventional GC-MS platform. Fecal samples with lower butyrate concentrations are measured more accurately on the rapid metabolomic screen resulting in a bias due to increased sensitivity compared to the GC-MS.

A negative bias is observed when comparing the QTRAP mass spectrometer measurement of DCA in the rapid metabolomic screen to our conventional measurements using a QToF. The rapid metabolomic screen employs the 3-NPH derivatization reagent with highly selective QTRAP detection of precursor to product ion transitions, which results in a very high signal-to-noise ratio. In contrast, the QToF analysis relies on a high mass accuracy to detect DCA by targeting the intact *m/z* without derivatization, which is less selective and results in greater levels of noise. Derivatization can improve sensitivity by reducing interference from otherwise coeluting compounds at the lower concentrations of DCA in the rapid screen.

The development of a rapid, sensitive and robust LC-MS/MS based screen for microbiota-derived metabolites represents a major improvement in the clinical assessment of the microbiome in patients. Although advances in genomics allow comprehensive investigation of the gut microbiome, sample preparation and analysis are lengthy and require significant laboratory and bioinformatic resources unavailable to most hospitals. With the increased use of LC-MS/MS technology in clinical chemistry laboratories, easily accessible reagents, and minimal sample preparation procedures, the rapid metabolic screen described herein can be readily implemented as a clinical test.

In summary, the unified LC-MS/MS-based metabolomic screen we have developed rapidly quantifies butyrate and DCA from fecal samples to measure gut microbiome functional dysbiosis. The assay has the flexibility to measure additional microbiome-relevant metabolites, such as other short-chain fatty acids and bile acids, which may be applicable to patients with specific diseases. The simulated fecal matrix presented here serves as an analyte-free surrogate for patient stool, enabling development, validation, and adherence to regulatory and accreditation standards (e.g. Clinical Laboratory Improvement Amendments, CLIA; College of American Pathologists, CAP) for clinical assays to assess health of the gut microbiome.

## ACKNOWLEDGEMENTS

We acknowledge the clinical team at the Duchossois Family Institute for consenting patients and collecting biological samples; the Host-Microbe Metabolomics Facility (HMMF), the Microbial Metagenomics Facility (MMF), the Bioinformatics team, and Translational Informatics team at the Duchossois Family Institute for technical assistance; and all members of the DFI for enthusiastic discussions that guided this study. We also acknowledge The Center for Research Informatics (CRI) at The University of Chicago for assistance with medical data extraction.

## Contributions

M.W.M., A.M., A.H., E.G.P., and A.M.S. were responsible for study conception and design. M.W.M., A.M., A.H., D.M., M.M., A.R., and J.L. collected mass spectrometry data. M.W.M., A.M., A.H., D.M., M.M., A.R., J.L. and A.M.S. performed data analysis and interpretation. A.B.N., B.K.P., C.J.L., M.A.O., and E.G.P. completed clinical studies. M.W.M., A.M., E.G.P., and A.M.S. drafted the article. All authors provided critical revision of the article and approved of the final version of the paper for publication.

## Notes

E.G.P. has filed a provisional patent application for the use of stool metabolite measurements as a method to identify patients at risk for hospital-acquired infection. E.G.P. is an inventor on patent numbers 11471495 (methods and composition for reducing infection), 11364269 (methods and composition for reducing infection or colonization), 11207374 (lantibiotics, lantibiotic-producing bacteria, compositions, and methods of production and use thereof), 11197897 (methods and compositions for reducing vancomycin-resistant Enterococci infection or colonization), 11045506 (methods and compositions for identifying and treating subjects at risk for checkpoint blockade therapy associated colitis) and receives royalties from Seres Therapeutics Inc. E.G.P. serves on the Scientific Advisory Board of Diversigen and on the Gates Foundation Global Health Scientific Advisory Committee. A.M.S. serves on the Scientific Advisory Board of 3^2^ Biosciences Gateway Biome.

## DATA AVAILABILITY

In compliance with the Health Insurance Portability and Accountability Act (HIPAA) and the University of Chicago institutional review board policy, clinical data obtained through the Center for Research Informatics (CRI) are stored on secure, encrypted and password-protected servers and are not all publicly available. Metagenomic information is publicly available on NCBI under BioProject numbers PRJNA912122 (liver disease patient cohort), PRJNA1039781 (liver transplant patient cohort), PRJNA845905 (heart transplant patient cohort), PRJNA1134172 (MICU patient cohort). Raw mass spectrometry data files will be made publicly available as a MassIVE repository upon peer review.

## REFERENCES

1. Michonneau, D. et al. Metabolomics analysis of human acute graft-versus-host disease reveals changes in host and microbiota-derived metabolites. Nat. Commun. 10, 5695 (2019).

2. Piccioni, A. et al. Gut Microbiota and : What We Know and the New Frontiers. Int. J. Mol. Sci. 23, (2022).

3. Odenwald, M. A. et al. Bifidobacteria metabolize lactulose to optimize gut metabolites and prevent systemic infection in patients with liver disease. Nat Microbiol 8, 2033–2049 (2023).

4. Lehmann, C. J. et al. Fecal metabolite profiling identifies liver transplant recipients at risk for postoperative infection. Cell Host Microbe 32, 117–130.e4 (2024).

5. Dela Cruz, M. et al. Reduced immunomodulatory metabolite concentrations in peri-transplant fecal samples from heart allograft recipients. Front Transplant 2, 1182534 (2023).

6. Stutz, M. R. et al. Immunomodulatory fecal metabolites are associated with mortality in COVID-19 patients with respiratory failure. Nat. Commun. 13, 6615 (2022).

7. Whon, T. W., Shin, N.-R., Kim, J. Y. & Roh, S. W. Omics in gut microbiome analysis. J Microbiol 59, 292–297 (2021).

8. Damhorst, G. L., Adelman, M. W., Woodworth, M. H. & Kraft, C. S. Current Capabilities of Gut Microbiome-Based Diagnostics and the Promise of Clinical Application. J. Infect. Dis. 223, S270–S275 (2021).

9. Wang, W.-L. et al. Application of metagenomics in the human gut microbiome. World J. Gastroenterol. 21, 803–814 (2015).

10. Fiori, J., Turroni, S., Candela, M. & Gotti, R. Assessment of gut microbiota fecal metabolites by chromatographic targeted approaches. J. Pharm. Biomed. Anal. 177, 112867 (2020).

11. Application of liquid chromatography–mass spectrometry to measure short-chain fatty acids in blood. Journal of Chromatography B 877, 719–724 (2009).

12. Liao, H.-Y. et al. Development of an Efficient and Sensitive Chemical Derivatization-Based LC–MS/MS Method for Quantifying Gut Microbiota-Derived Metabolites in Human Plasma and Its Application in Studying Cardiovascular Disease. J. Proteome Res. (2021) doi:10.1021/acs.jproteome.1c00147.

13. Clinical Laboratory Improvement Amendments (CLIA). Standards and Certification: Laboratory Requirements vol. 42 CFR 493 678–812 (1988).

14. Haak, B. W. et al. Impact of gut colonization with butyrate-producing microbiota on respiratory viral infection following allo-HCT. Blood 131, 2978–2986 (2018).

