## Supplemental Information for "Fecal Metabolite Quantitation for Rapid Assessment of the Gut Microbiome"

**Authors Info & Affiliations**:

[1]Duchossois Family Institute, University of Chicago, Chicago, IL, 60637, USA

[2]Department of Pathology, University of Chicago, Chicago, IL, 60637, USA

[3]Department of Medicine, University of Chicago, Chicago, IL, 60637, USA

[4]Department of Microbiology, University of Chicago, Chicago, IL, 60637, USA

**Table S1** Sample preparation and acquisition time-frame in minutes to acquire quantitative concentration values for one biological sample using conventional metabolomics methods and the rapid metabolomic screen.

| MS Platform | dry down | derivatization | additional [a] | acquisition [b] | analysis |
|---|---|---|---|---|---|
| QToF LC-MS | 120 | 0 | 60 | 359 | 10 |
| PFB-Br GC-ECNI-MS | 0 | 60 | 20 | 228 | 10 |
| Rapid Screen | 0 | 30 | 10 | 160 | 10 |

[a] Additional sample preparation for QTof LC-MS method involves resuspension, centrifugation, and transfer to a new MS vial, additional steps in PFB-Br GC-ECNI-MS involve sample cooling, centrifugation, and transfer to a new MS vial, while additional steps in the rapid metabolomic screen involve sample handling and reaction quenching at -80 °C; [b] Acquisition time includes the blanks, quality control, and calibration curve sample injections required to generate reliable quantification of metabolites in a single biological sample.

**Table S2** Data acquisition parameters for the Sciex QTRAP used in the rapid metabolomic screen.

| Metabolite | Precursor ion (Q1 m/z; Da) | Quantifier product ion (Q3 m/z; Da) | Qualifier product ion (Q3 m/z; Da) | Retention time (min) | Acquisition window (sec) | Dwell Weight | Collision Energy |
|---|---|---|---|---|---|---|---|
| 3NPH-isobutyrate | 222.09 | 137.1 | 152.0 | 2.2 | 30 | 1 | -30 |
| 3NPH-butyrate | 222.09 | 137.1 | 152.0 | 2.5 | 30 | 1 | -30 |
| 3NPH-UDCA[a] | 526.33 | 152.0 | 137.1 | 5.1 | 30 | 1 | -65 |
| 3NPH-HDCA[b] | 526.33 | 152.0 | 137.1 | 5.2 | 30 | 1 | -65 |
| NPH-isoDCA[c] | 526.33 | 152.0 | 137.1 | 5.6 | 30 | 1 | -65 |
| NPH-CDCA[d] | 526.33 | 152.0 | 137.1 | 6.2 | 30 | 1 | -65 |
| NPH-DCA[e] | 526.33 | 152.0 | 137.1 | 6.8 | 30 | 1 | -65 |
| 3NPH-3-DCA[f] | 526.33 | 152.0 | 137.1 | 7.4 | 30 | 1 | -65 |
| 3NPH-$D_7$-butyrate | 229.09 | 137.1 | 152.0 | 2.5 | 30 | 1 | -30 |
| 3NPH-$D_4$-CDCA[d] | 530.33 | 152.0 | 137.1 | 6.8 | 30 | 1 | -65 |
| 3NPH-$D_4$-DCA[e] | 530.33 | 152.0 | 137.1 | 6.2 | 30 | 1 | -65 |

[a] UDCA, ursodeoxycholic acid; [b] HDCA, hyodeoxycholic acid; [c] isoDCA, isodeoxycholic acid; [d] CDCA, chenodeoxycholic acid; [e] DCA, deoxycholic acid; [f] 3-DCA, 3-deoxycholic acid.

**Table S3** Summary of performance metrics for rapid metabolite screen.

| Performance Characteristic | Butyrate | Deoxycholic acid |
|---|---|---|
| **Analytical Measurement Range** | 4.3 µM–3030.1 µM | 0.9 µM–64.9 µM |
| **Lower Limit of Quantitation** | 9.75 µM | 0.9 µM |
| **Inter-Day Precision** (%CV, n = 14 across 8 runs) | 2.43–10.4% | 7.4–14.3% |
| **Recovery** (across 5 concentrations, n = 14) | 98.2–109% | 92.4–109.7% |
| **Isomers Accounted For** | isobutyrate | UDCA, HDCA, 3-DCA, CDCA, isoDCA |

**Table S4** Comparison of butyrate and deoxycholic acid concentration between the rapid metabolomic screen and conventional mass spectrometry-based methods with patient clinical metadata. 'Greater-than' or 'less-than' values indicate metabolite concentrations above and below rapid screen AMR and/or LLOQ.

| | Clinical Metadata | | | Butyrate (µM) | | Deoxycholic acid (µM) | |
|---|---|---|---|---|---|---|---|
| **Deidentified Patient Sample ID** | **Patient cohort** | **Race** | **Sex** | **Rapid Screen** | **PFB-Br - GCMS** | **Rapid Screen** | **Q-ToF LCMS** |
| HT_009_03 | Heart transplant | Black or African-American | Male | 545.36 | < 750 | 1.82 | < 2.50 |
| HT_027_03 | Heart transplant | White | Male | 2626.80 | 3460 | > 64.90 | 523.43 |
| HT_032_18 | Heart transplant | White | Male | < 9.75 | < 750 | < 0.90 | < 2.50 |
| HT_042_07 | Heart transplant | Black or African-American | Female | < 9.75 | < 750 | < 0.90 | < 2.50 |
| HT_048_01 | Heart transplant | Black or African-American | Male | 145.76 | < 750 | < 0.90 | < 2.50 |
| HT_061_02 | Heart transplant | Black or African-American | Male | < 9.75 | < 750 | < 0.90 | < 2.50 |
| HT_066_06 | Heart transplant | Black or African-American | Female | 57.23 | < 750 | 1.86 | < 2.50 |
| HT_071_06 | Heart transplant | Asian/Mideast Indian | Female | 82.24 | < 750 | < 0.90 | < 2.50 |
| HT_080_04 | Heart transplant | Black or African-American | Female | < 9.75 | < 750 | < 0.90 | < 2.50 |

| | Clinical Metadata | | | Butyrate (µM) | | Deoxycholic acid (µM) | |
|---|---|---|---|---|---|---|---|
| **Deidentified Patient Sample ID** | **Patient cohort** | **Race** | **Sex** | **Rapid Screen** | **PFB-Br - GCMS** | **Rapid Screen** | **Q-ToF LCMS** |
| HT_106_01 | Heart transplant | Black or African-American | Male | > 3030.10 | 3440 | > 64.90 | 1606.54 |
| LD_001_01 | Liver disease | Black or African-American | Male | 1190.46 | 1660 | 8.22 | 4.87 |
| LD_011_01 | Liver disease | White | Female | > 3030.10 | 3570 | 30.38 | 12.07 |
| LD_014_02 | Liver disease | White | Male | 2866.56 | 3090 | > 64.90 | 20.62 |
| LD_023_01 | Liver disease | More than one Race | Male | 2594.94 | 2860 | 25.75 | 8.87 |
| LD_025_03 | Liver disease | Unknown or Patient unable to respond | Female | < 9.75 | < 750 | < 0.90 | < 2.50 |
| LD_046_01 | Liver disease | Black or African-American | Female | 2076.40 | 2580 | 15.84 | 14.90 |
| LD_049_01 | Liver disease | White | Male | 239.53 | 840 | 16.21 | 14.57 |
| LD_061_01 | Liver disease | White | Female | 2433.77 | 2970 | 35.44 | 27.49 |
| LD_070_01 | Liver disease | Black or African-American | Female | 1439.95 | 1950 | 5.12 | 4.25 |
| LD_087_01 | Liver disease | Black or African-American | Female | 1000.91 | 920 | 3.82 | 4.38 |
| LD_095_01 | Liver disease | White | Female | < 9.75 | < 750 | < 0.90 | < 2.50 |
| LD_129_27 | Liver disease | Black or African-American | Female | 84.56 | < 750 | < 0.90 | < 2.50 |
| LD_134_04 | Liver disease | White | Male | < 9.75 | < 750 | < 0.90 | < 2.50 |
| LD_136_08 | Liver disease | White | Male | > 3030.10 | 14960 | < 0.90 | 2.85 |
| LD_155_01 | Liver disease | Black or African-American | Male | < 9.75 | < 750 | < 0.90 | < 2.50 |
| LD_156_01 | Liver disease | Black or African-American | Male | 31.21 | < 750 | 2.68 | 3.21 |
| LD_159_01 | Liver disease | More than one Race | Female | 563.70 | < 750 | 3.68 | 3.19 |

| | Clinical Metadata | | | Butyrate (µM) | | Deoxycholic acid (µM) | |
|---|---|---|---|---|---|---|---|
| **Deidentified Patient Sample ID** | **Patient cohort** | **Race** | **Sex** | **Rapid Screen** | **PFB-Br - GCMS** | **Rapid Screen** | **Q-ToF LCMS** |
| LD_174_01 | Liver disease | Black or African-American | Female | < 9.75 | < 750 | < 0.90 | < 2.50 |
| LD_181_01 | Liver disease | Black or African-American | Male | 1199.18 | 1670 | 4.93 | 5.66 |
| LD_191_01 | Liver disease | More than one Race | Male | 2787.93 | 3430 | 36.49 | 38.23 |
| LD_192_01 | Liver disease | Black or African-American | Male | > 3030.10 | 3610 | < 0.90 | 2.53 |
| LD_193_01 | Liver disease | White | Female | 1908.16 | 2420 | 35.35 | 35.81 |
| LD_194_08 | Liver disease | White | Male | > 3030.10 | 8340 | < 0.90 | < 2.50 |
| LD_195_01 | Liver disease | White | Male | 2899.99 | 3390 | 61.26 | 54.81 |
| LD_228_01 | Liver disease | Black or African-American | Female | 362.53 | 1010 | 1.14 | < 2.50 |
| LD_234_11 | Liver disease | Asian/Mideast Indian | Female | < 9.75 | < 750 | 1.73 | < 2.50 |
| LD_242_01 | Liver disease | More than one Race | Female | 1970.14 | 2430 | 5.55 | < 2.50 |
| LD_262_01 | Liver disease | More than one Race | Male | 225.06 | 810 | 15.51 | 7.72 |
| LD_264_06 | Liver disease | White | Male | 1789.05 | 2440 | > 64.90 | 143.37 |
| LD_284_02 | Liver disease | Patient declines to respond | Female | > 3030.10 | 4000 | < 0.90 | < 2.50 |
| LD_293_01 | Liver disease | White | Male | 2208.65 | 2560 | 17.91 | 7.32 |
| LD_302_01 | Liver disease | White | Male | 41.27 | < 750 | < 0.90 | < 2.50 |
| LD_304_01 | Liver disease | White | Female | 2360.53 | 2430 | 12.09 | 7.69 |
| LD_313_01 | Liver disease | Black or African-American | Female | 27.67 | < 750 | 3.21 | < 2.50 |
| LD_314_01 | Liver disease | Black or African-American | Female | 1458.83 | 1960 | < 0.90 | 3.36 |
| LD_322_11 | Liver disease | White | Female | < 9.75 | < 750 | < 0.90 | < 2.50 |
| LD_338_02 | Liver disease | More than one Race | Female | 629.40 | 820 | 26.35 | 32.43 |

| | Clinical Metadata | | | Butyrate (µM) | | Deoxycholic acid (µM) | |
|---|---|---|---|---|---|---|---|
| **Deidentified Patient Sample ID** | **Patient cohort** | **Race** | **Sex** | **Rapid Screen** | **PFB-Br - GCMS** | **Rapid Screen** | **Q-ToF LCMS** |
| LD_347_01 | Liver disease | Black or African-American | Female | < 9.75 | < 750 | < 0.90 | 2.85 |
| LD_394_01 | Liver disease | Black or African-American | Female | 2018.87 | 2350 | 32.24 | 43.53 |
| LD_411_02 | Liver disease | Black or African-American | Female | 281.17 | < 750 | < 0.90 | < 2.50 |
| LD_413_01 | Liver disease | White | Male | 2263.81 | 2540 | 7.03 | 6.44 |
| LD_414_03 | Liver disease | Black or African-American | Male | 183.40 | < 750 | < 0.90 | < 2.50 |
| LD_446_01 | Liver disease | Unknown or Patient unable to respond | Female | 2952.59 | 3280 | > 64.90 | 62.08 |
| LD_470_01 | Liver disease | White | Male | 1010.40 | 1310 | 8.81 | 10.44 |
| LD_481_01 | Liver disease | White | Female | 300.33 | < 750 | 7.69 | 7.67 |
| LD_494_01 | Liver disease | Asian/Mideast Indian | Female | 507.24 | < 750 | 10.28 | 13.69 |
| MICU_063_02 | Medical intensive care unit | White | Male | > 3030.10 | 2350 | > 64.90 | 143.72 |
| MICU_082_03 | Medical intensive care unit | Black or African-American | Male | 2352.20 | 2180 | 1.38 | < 2.50 |
| MICU_103_01 | Medical intensive care unit | Black or African-American | Female | 69.63 | 250 | 16.11 | 2.67 |
| MICU_107_03 | Medical intensive care unit | Black or African-American | Female | 24.46 | < 750 | 7.72 | 2.73 |
| MICU_118_06 | Medical intensive care unit | White | Male | 257.55 | < 750 | 12.92 | 7.57 |
| MICU_181_01 | Medical intensive care unit | Black or African-American | Male | > 3030.10 | 3860 | > 64.90 | 106.88 |
| MICU_187_01 | Medical intensive care unit | Black or African-American | Female | 188.92 | < 750 | > 64.90 | 58.38 |
| MICU_199_05 | Medical intensive care unit | Black or African-American | Male | < 9.75 | < 750 | 17.72 | 10.34 |

| | Clinical Metadata | | | Butyrate (µM) | | Deoxycholic acid (µM) | |
|---|---|---|---|---|---|---|---|
| **Deidentified Patient Sample ID** | **Patient cohort** | **Race** | **Sex** | **Rapid Screen** | **PFB-Br - GCMS** | **Rapid Screen** | **Q-ToF LCMS** |
| MICU_205_01 | Medical intensive care unit | White | Male | 941.73 | 1110 | < 0.90 | < 2.50 |
| MICU_238_02 | Medical intensive care unit | White | Male | 1268.99 | < 750 | 5.59 | 3.63 |
| MICU_260_03 | Medical intensive care unit | White | Male | 1742.45 | 1960 | > 64.90 | 106.78 |
| MICU_318_02 | Medical intensive care unit | White | Male | < 9.75 | < 750 | 9.80 | 8.23 |
| MICU_331_04 | Medical intensive care unit | Asian/Mideast Indian | Female | 99.90 | < 750 | 14.37 | 3.92 |
| MICU_367_03 | Medical intensive care unit | White | Female | < 9.75 | < 750 | < 0.90 | < 2.50 |
| MICU_426_02 | Medical intensive care unit | Black or African-American | Female | 2028.07 | 2600 | > 64.90 | 133.74 |
| MICU_436_04 | Medical intensive care unit | Black or African-American | Male | < 9.75 | < 750 | < 0.90 | < 2.50 |
| MICU_494_01 | Medical intensive care unit | White | Female | 1568.62 | 1600 | < 0.90 | < 2.50 |
| UC_021_07 | Liver transplant | White | Female | 1810.27 | 1130 | > 64.90 | 278.32 |
| UC_050_02 | Liver transplant | White | Female | < 9.75 | < 750 | < 0.90 | < 2.50 |
| UC_077_04 | Liver transplant | White | Female | < 9.75 | 860 | < 0.90 | < 2.50 |
| UC_136_01 | Liver transplant | Black or African-American | Male | 2804.84 | 3820 | > 64.90 | 64.27 |
| UC_149_01 | Liver transplant | Black or African-American | Male | > 3030.10 | 3780 | > 64.90 | 1111.21 |
